# Assessing the origins of the European Plagues following the Black Death: a synthesis of genomic, historical and ecological information

**DOI:** 10.1101/2021.04.20.440561

**Authors:** Barbara Bramanti, Yarong Wu, Ruifu Yang, Yujun Cui, Nils Chr. Stenseth

**Affiliations:** Centre for Ecological and Evolutionary Synthesis (CEES), Department of Biosciences, University of Oslo, Blindern, N-0316, Oslo, Norway; Department of Neuroscience and Rehabilitation, Faculty of Medicine, Pharmacy and Prevention, University of Ferrara, 44121 Ferrara, Italy; State Key Laboratory of Pathogen and Biosecurity, Beijing Institute of Microbiology and Epidemiology, Beijing, 100071, China; Ministry of Education Key Laboratory for Earth System Modeling, Department of Earth System Science, Tsinghua University, Beijing, 100084, China

**Keywords:** *Yersinia pestis*, ancient DNA, Black Death, European Plague, pathogen evolution

## Abstract

The Second Plague Pandemic started in Europe with the Black Death in 1346 and lasted until the 19^th^ century. Based on ancient DNA studies, there is a scientific disagreement over whether the bacterium, *Yersinia pestis*, came into Europe once (Hypothesis 1), or repeatedly over the following four centuries (Hypothesis 2). Here we synthesize the most updated phylogeny together with historical, archeological, evolutionary and ecological information. On the basis of this holistic view, we conclude that Hypothesis 2 is the most plausible. We also suggest that *Y. pestis* lineages might have developed attenuated virulence during transmission, which can explain the convergent evolutionary signals, including *pla*-decay, that appeared at the end of the pandemics.

**Significance Statement:** Over the last few years there has been a great deal of scientific debate regarding whether the plague bacterium, *Yersinia pestis*, spread from a Western European reservoir during the Second Plague Pandemic, or if it repeatedly came to Europe from Asia. Here we make a synthesis of the available evidence, including genomes of ancient DNA, historical, archeological and ecological information. We conclude that the bacterium most likely came to Europe from Asia several times during the Second Plague Pandemic.

## Main Text

Researchers agree that the Second Plague Pandemic was caused by *Yersinia pestis* (1-9), which arrived in Europe from Caffa transported by Genoese galleys on the Black Sea at the beginning of the Black Death (10). However, there is no consensus among researchers as to the origins of plague epidemics in Europe following the Black Death and ravaging Europe until the 19^th^ century, as attested by historical documents (11).

The two main theories are that one or more plague reservoirs remained in Western Europe during the entire Second Plague Pandemic (referred to in the following as Hypothesis 1) (3, 4, 8, 12), or the bacteria repeatedly invaded Europe from non-Western European reservoir(s) during the same period (referred to in the following as Hypothesis 2) (6, 7, 9, 11, 13). Here, we assess these two hypotheses using a broad spectrum of evidence including historical and archeological, genetic and evolutionary, as well as ecological information.

## Results and Discussion

### Assessment of the two hypotheses

For the purpose of understanding the evolution of the plague bacteria, more than 100 ancient *Y. pestis* genomes have been published to date. The last 17 were recently reported during a short period by four distinct research groups (7-9, 12). Using most of the ancient genomes (criteria for exclusion are described in methods) along with 499 modern ones, we present here the most updated phylogeny (Fig. 1).

**Fig 1.**
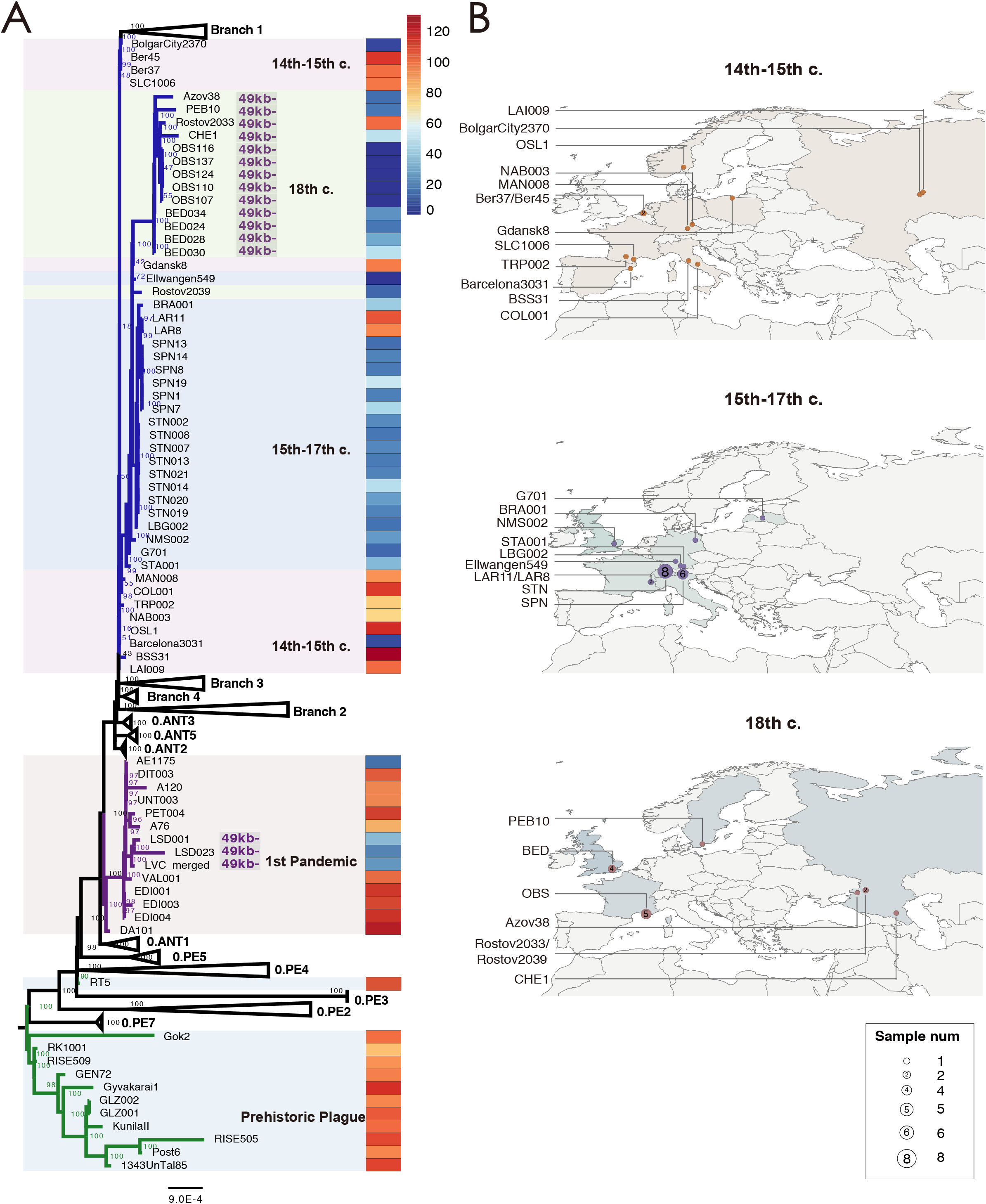
Phylogeny and archaeological site locations of ancient genomes. (A) A maximum likelihood phylogeny was obtained with 574 genomes of *Y. pestis* (including 75 ancient genomes) involved, based on 12,608 SNPs. The numbers at each node indicate the bootstrap values of 1,000 replicates. Branches highlighted in blue correspond to the Second Pandemic, which is subdivided in three groups: the 14^th^-15^th^ century group, which also includes the Black Death and the *Pestis secunda* (1357-1366) strains, the 15^th^-17^th^ century group, and the 18^th^ century group (which includes also the BED genomes for homogeneity). Branches in purple correspond to the First Pandemic and branches in green correspond to the prehistoric plague. The ratio between the depth of *pla* and that of the entire pPCP1 plasmid for all ancient genomes is shown in the rightmost heatmap, with a color scale ranging from 0 (dark blue) to 130+ (dark red). (B) Geographic distribution of the three waves during the Second Pandemic.

The updated phylogeny confirms the almost clonal nature of the Black Death strains, in comparison to all other lineages of the Second Plague Pandemic, including the strains from the *Pestis secunda* (Ber37 and Ber 45, The Netherlands (6), and BolgarCity2370, Russia (3)), which are placed on Branch 1 (see also London-Ind6330, UK (3)), as well as to all other strains, which are placed on the post-Black Death branch. There is general agreement that the post-Black Death branch was hosted in a novel wild rodent reservoir – either in Europe or outside Europe. The original hypothesis (Hypothesis 1) claims that such a plague reservoir existed in Western Europe (14), perhaps in the Alps (15). However, a newer hypothesis (Hypothesis 2) claims that the plague reservoir was in Asia, possibly close to Eastern Europe (6, 7, 9, 11, 13).

In order to more easily view the phylogeny from the Second Plague Pandemic and to better contrast the evidence for the two hypotheses, we generated two schematic figures (Fig. 2) and a table (Table 1).

**Table 1.** Main differences between the two competing hypotheses proposed to explain the phylogeny of *Y. pestis* of the Second Plague Pandemic. Genomic and evolutionary, historical and archaeological, as well as ecological arguments are considered.

|  | Hypothesis 1 | Hypothesis 2 |
| --- | --- | --- |
| Main differences |  |  |
| Origin of the outbreaks | Plague established in Western European reservoirs (for example, marmots in the Alps) (3, 4, 8) | Plague was repeatedly imported from Eastern European or Asian reservoirs (6, 7, 9, 11, 13) |
| Transmission | Mediated by rats infected by wild rodents, as in China during the Third Pandemic | Imported by rats, humans, and goods, and subsequently spread by chains of human transmission, as in Europe during the Third Pandemic (16, 17) |
| Vector | *Xenopsylla cheopis* and other rodent fleas | Any ectoparasite, including *Pulex irritans*, and body lice (18, 19) |
| Supporting information |  |  |
| Population genomics | Western European strains (the Alpine clade) are basal in the sub-branch (although of different eras). A model proposes Western European strains as ancestral sources for the transmissions (Fig. S1) | The oldest (LAI009) and the most recent strains, in addition to Bolgar, are all from Eastern Europe (Russia), as well as all the strains of the 18<sup>th</sup> century; the majority of the Western European strains in the phylogeny come from ports |
| History and archaeology | A hypothesis (15) suggests that the plagues from the 16<sup>th</sup> century in the Alps, were not imported by major trade centers | Multiple historical records assert that plague was imported for outbreaks associated with the Black Death strains (partially reviewed in (6, 7, 11, 13)), for BRA (4), for SPN (in the Alps), PEB10 (7) and OBS (5). Multiple records of importation are historically attested (20), particularly in harbors |
| Climate | No climatic signal in support, although 4 datasets of climatic proxies were from the Alps (13) | Strong signals of climate-driven introductions of plague from Asia (13) |
| Evolution | *Y. pestis* developed in Western European reservoir strains *pla*<sup>+</sup>/*pla*<sup>-</sup> and Delta49kb possibly as a form of adaptation to the local host (rodents). | *Y. pestis* developed *pla*<sup>+</sup>/*pla*<sup>-</sup> and Delta49kb strains possibly as a form of adaptation to the new host (humans) and/or new vectors (fleas or body lice). |

**Fig. 2.**
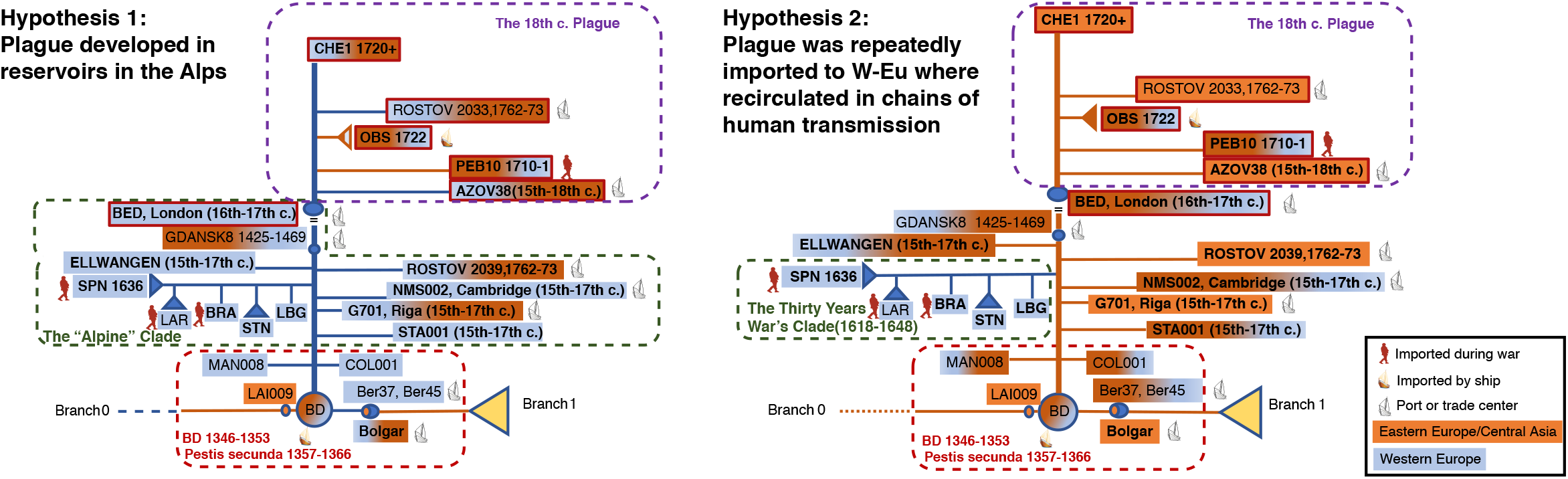
Schematic comparison between the two main hypotheses for the interpretation of the *Y. pestis* phylogeny of the Second Plague Pandemic. Historic and evolutionary information is included in the schematic figures. In addition to the symbols explained in the figure, we outlined in red the strains showing the 49kb deletion. *Pla*-decay (meaning both, full or partial absence of the *pla*-gene) is indicated by the names in bold.

Hypothesis 1 is supported by a phylogenetic analysis based on the currently available ancient genomes, which infers high posterior probability for a Western European source of the transmissions on the post Black Death branch (see Fig. S1). However, as the dataset includes 41 ancient genomes from Western Europe against only 8 strains from Eastern Europe (including Gdansk and Riga), the proposed origins from Western Europe are likely to be biased toward a European reservoir due to a size-effect bias. Notably, the most basal genome LAI009 (4) (the Black Death’s lineage), Bolgar (at the root of Branch 1), and the most recent genome (CHE1 (7)), all originated from Western Russia, implying that they might have been closer to a putative Asian or Eastern European reservoir. This continuity does represent strong evidence in support of Hypothesis 2.

Using only genomic data, Hypothesis 1 might be seen as the most parsimonious hypothesis, since it proposes an internal source for all western Eurasian outbreaks. However, for two locations (Pestbacken, Sweden 1710 (PEB10), and Marseille, France 1722 (OBS)), an origin from the Ottoman Empire is historically and archaeologically well supported (7). Thus, Hypothesis 1 needs to account for a back- and-forth spread, which reintroduced plague on two occasions to the Ottoman Empire and back again to Western Europe. Notably, none of the strains from the 18^th^ century appear to have originated in Western Europe according to historical sources (7, 9).

Hypothesis 1 assumes the existence of a wild rodent plague reservoir in the Alps, which is not supported by ecological evidence (13). Instead, a study of more than 7,000 historical plague outbreaks and 15 tree-ring datasets (4 of which from the Alps), found climatic signals in support of frequent re-importations of plague from Asia into Eastern and Western European harbors (13).

Intriguingly, only a few genotyped strains are nodes on the backbone of the post-Black Death branch: the strains of the Black Death itself, the strain from Gdansk 1425-1469 and the strains from London (BED, 16^th^-17^th^ century). While the strains of the Black Death were notoriously imported into Western Europe from the Mongol Empire via Caffa in Crimea (10), both Gdansk and London were very active harbors also in historical times and were very often hit by plague. Interestingly, *Yersinia pestis* was also recovered from a rat found in Gdansk. Although the genome is partial, due to the different SNPs profile, it is clear that the strain from the rat could not have infected the victim (Gdansk8) (9). Being a port, Gdansk may indeed have hosted diverse importations of infected rats in the period 1425-1469, as it happened in European harbors during the Third Pandemic (16).

Hypothesis 2 is consistent with the ecological as well as with the historical evidence (Fig. 2 and Table 1). The only Western European sub-cluster, the ‘Alpine cluster’ formed by LBG, STN, BRA, LAR and SPN, may naturally be explained by the circulation of soldiers and troops in Europe during the Thirty Years War (1618-1648), which made up human chains of transmission with historically documented epidemic events (20, 21). For three strains, SPN from the Italian Alps, LAR from the French Alps and BRA, from Northern Germany, the relationship with the time of the Thirty Years War is historically and archaeologically documented (4, 7, 12). Human chains of transmission, which do not require the presence of rats to start and sustain an epidemic, might explain the circulation of the plague within Europe over long periods of time. They might be due to interpersonal contacts, crowding, infected parasites in clothes or goods, or contact with infected pets. Several chains of human transmission within Europe could be reconstructed for cases of the last century (16, 17).

### Two convergent mutations

To better understand the evolution of *Y. pestis*, we examined two more mutations, which were recently discovered in ancient strains. In the most recent sub-clade of the Second Pandemic, starting with BED, there is a 49kbp deletion with unknown function. This deletion was also present in the last lineage of the First Pandemic, and, in both cases, might have accounted for the decline of the pandemic (7). We found the same mutation in the Rostov 2033 strain in the 18^th^ century clade (Fig.1 and 2). By contrast, a second strain found in the same cemetery in Rostov (Rostov 2039) has a different SNPs-pattern and lacks the chromosomal deletion.

Another mutation, the depletion of the *pla*-gene on the plasmid pPCP1, has recently been proposed as the cause of the disappearance of the Second Plague Pandemic in the 18^th^ century (8), given that the *pla*-gene is an important virulence factor of *Y. pestis*. We checked for the presence of the *pla*+/*pla*-plasmids in all published ancient strains. The ratio in coverage depth between *pla* and the whole pPCP1 plasmid indicates the status of *pla*-loss in an organism (Fig. 3). If the depth of *pla* is significantly lower than that of pPCP1, it might properly be concluded that the *pla*-gene was lost in some pPCP1 plasmids. Our analyses show that the ratio of *pla* in the Black Death and post-Black Death genomes appears to be different when compared with the pre-historic and the First pandemic lineages (Fig. 1). We have also checked randomly selected modern *Y. pestis* genomes in different lineages, and their depth of *pla* and pPCP1 are quite consistent, indicating no other *pla*-loss in modern plagues. By contrast, the generalized depletion of *pla* extensively observed during the post-Black Death era and at the end of the First Pandemic (Fig. 1) seems to be consistent. Given that the sequencing data were generated by several different research groups, a systemic error during sequencing is unlikely.

**Fig 3.**
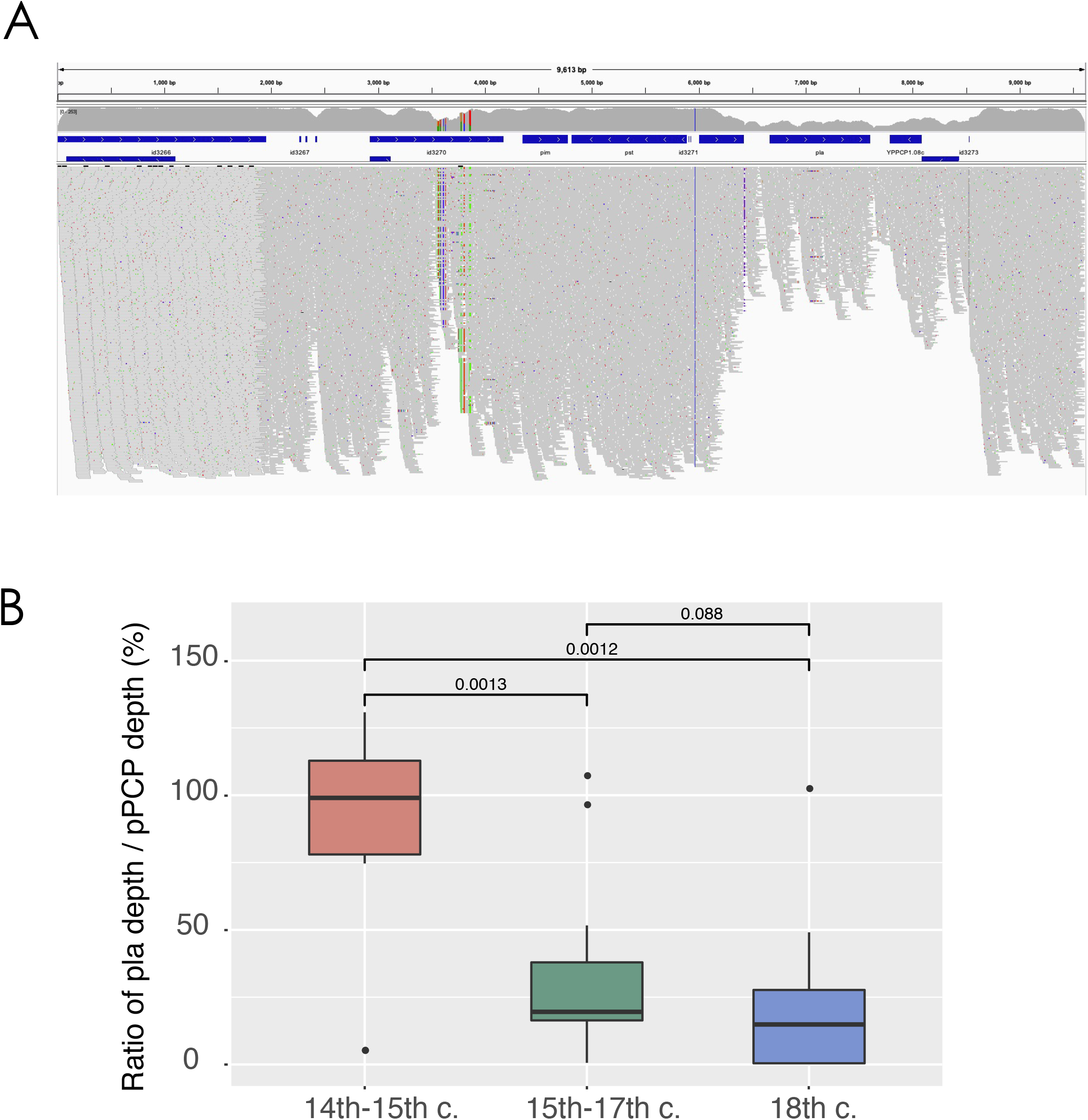
The decay of the *pla*-gene. (A) Depth plot of the pPCP1 plasmid in strain CHE1 using IGV. The annotated genes of the pPCP1 plasmid are marked with blue bars. The average sequencing depth of whole pPCP1 plasmid is 195.65X, while the average sequencing depth of the *pla*-region is 96.04X. (B) Group-wise comparison of the ratio between the depth of *pla* and that of whole pPCP1 plasmid among three waves of the Second Pandemic. Boxplots depict the upper, median, and lower quartiles of the ratios, individual dots indicate outliers that lie outside of 1.5 times the interquartile range, and vertical lines indicate the range of all ratios except for outliers. The *p*-value of group-wise comparison using the Wilcoxon-test are labeled on the top, two of which are statistically significant (*p*<0.05). Data of Fig. 3b are provided in Dataset S4.

It seems that full *pla*-strains were slightly depleted at the end of the Second Pandemic (8), with the same phenomenon at the end of the First Pandemic. Notably, however, Rostov2033, one of the most recent genomes of the Second Pandemic, shows full reads pPCP1 plasmids, whereas CHE, the most recent historical strain, shows very slight *pla*-decay (Fig. 3). This observation is not fully in agreement with the proposed hypothesis that *pla*-depletion contributed to the end of the pandemic. An alternative explanation for this phenomenon (8) is that the differences observed in the full *pla*-plasmids might be due to different forms of plague. In particular, bubonic plague and pneumonic plague need the *pla*-gene to develop, whereas primary septicemic plague does not (8). It seems that plague existed in all three forms, at least from the time of the First Pandemic, however, this does not add any specific evolutionary information to the observed variability.

### The evolution of the *pla*-gene

We propose an evolutionary hypothesis for the presence of lineages with *pla* decay. One of the optimized survival strategies for an emerging pathogen is to balance its virulence to the main host with its transmission strategy. This trade-off hypothesis was previously demonstrated for *Y. pestis* (22, 23). This mechanism would allow the bacterium to reduce virulence and enhance the time of survival of the host and, consequently, of the pathogen (24). After experiencing the Black Death and successive waves, the *pla* decay strains might have attempted to acquire a fitness advantage reducing their virulence by increasing the time-to-death. Indeed, we observe among the victims only *pla*+/*pla*-mixed strains, whereas *pla-*lineages might have survived longer in the host population, providing a milder form of the illness.

The Eastern European/Asia clade of the 18^th^ century (including CHE1) further lost the 49kb region, which can be the result of an extension of a virulence attenuated pattern. Such events of attenuated virulence might have occurred multiple times in the *Y. pestis* evolutionary history, and left out host-adapted lineages, such as for 0.PE2 and 0.PE4 (25). Therefore, the possible virulence reduction caused by *pla* decay and loss of the 49k region is not necessarily the reason for the extinction of plague at the end of the First and Second Pandemics, but might be the result of a form of adaptation to a new host, which may be the wild rodent in the putative Western European reservoir (Hypothesis 1), a new host in the Asian reservoir or the human host (Hypothesis 2), as well as their vectors. We observed that the newly published strains from Lariey (French Alps, (12)) do not show *pla*-decay, in contrast to other Alpine lineages (SPN). This evidence might exclude the hypothesis of an adaptation to a host in a Western European reservoir. Thus, we tentatively propose that this mechanism of *pla*-decay would support the presence of human-to-human transmission chains mediated by human ectoparasites (fleas and body lice) during plague pandemics in Europe, the plausibility of which has previously been demonstrated (16-18).

## Conclusion

Altogether, the most consistent interpretation of the current information is in support of Hypothesis 2. This implies that there must have existed a reservoir outside of Western Europe (with the Ottoman Empire, Persia and Central Asia as possible candidates (7, 9)). Such a reservoir could then fed, with multiple introductions, the Second Plague Pandemic outbreaks in Western Europe along Northern and Southern trade routes (6, 26). In addition there might have been continuous recirculation of plague in (Western and Eastern) Europe with the movement of goods and troops. The recirculation of plague within Western Europe, mediated by trade and travel, might mimic the presence of a local reservoir, assuring the long-lasting re-emergence of epidemics on the continent. Such recirculation of plague could explain the collateral branches on the phylogenetic tree – an example of which may be the Thirty Years War Clade (1618-1648) evidenced in Fig. 2. Additional ancient strains from Asia and Eastern Europe, and more accurate dating and historical contextualization, will provide further evidence to clarify the phylogeny and evolution of the *Y. pestis* pathogen.

## Methods

### SNP calling and evaluation

All raw reads of 111 published ancient genomes were downloaded from NCBI SRA and EBI ENA databases. For the clones from each sampling site, only one genome with the highest quality (sequencing depth) as reported in corresponding publications was chosen,and genomes covering less than 20% of the chromosome length of the CO92 assembly (GCF_000009065.1) were excluded from further analysis, which resulted in a dataset with a total of 75 ancient genomes (Dataset S1). We trimmed and quality filtered raw reads using Trimmomatic v0.38 (27), and reads shorter than 30 bp and below a quality score of 20 were discarded. Subsequently, the filtered reads were mapped against the CO92 assembly with BWA mem model (v0.7.17) (28) and the aligned reads were extracted from bam files using SAMtools (v1.9) (29) view command (-bF 4) and then different runs of the same sample were merged using SAMtools merge command. Sequences with more than 10 soft and hard clipped alignments were filtered out by samclip, and duplicates were removed using Picard’s MarkDuplicates module. SNP calling was performed using the UnifiedGenotyper of the Genome Analysis Toolkit (GATK v3.8) (30) under the “EMIT_ALL_CONFIDENT_SITES” option with a minimum confidence threshold 10; a vcf file for every ancient genome was produced and SNPs that were close to each other by less than 20 bp were excluded.

A total of 499 modern genomic assemblies of *Y. pestis* available in NCBI Genbank database on 19^th^ October, 2020 were downloaded (Dataset S2) and then aligned to the CO92 assembly using NASP’s convert_external_genome command (31), which was based on MUMmer’s nucmer and delta-filter modules (v3.23) (32). A fasta file 1-to-1 position aligned with a reference fasta for every modern genome was created.

All vcf files and aligned fasta files were used to aggregate sample calls into matrices with NASP’s matrix module. For ancient genomes, a SNP would be called when supported by >=3 reads and >= 90% allele frequency. After manually validating the calls for ancient genomes with notably longer branch lengths (SPN strains, BSS31, and SLC1006) according to their published SNP lists, we got a final dataset of 12,608 polymorphic loci (Dataset S3).

### Phylogenetic analyses and geographic extent of sampled sites

A fasta file, concatenated of all SNP sites, was used to generate a maximum-likelihood tree with 1,000 fast bootstrap replicates using IQ-TREE (v1.6.5) (33)with the option -m MFP+ASC to infer the best substitution model and account for ascertainment bias correction. Then FigTree (v1.4.3) was used to visualize the generated tree. The packages of ggplot2, maptoos and maps in R (v3.6.1) were used to mark the archaeological site locations of samples from the Second Plague Pandemic. The longitude and latitude for each site was taken from the website mapcoordinates (https://www.mapcoordinates.net/en).

### Phylognetic analyses with calculation of MCMC posterior probability

A maximum-likelihood tree for 47 ancient genomes during the Second Pandemic was rebuilt using RAxML(v8.2.11) (34) with 100 replicates and GTRGAMMA model. It was rerooted to strain LAI009 and transformed into newic format using FigTree. We used the ReorderData function in evobiR (v1.1, R package) to match the source records (country names) to the order of tips on the phylogenetic tree. Then the make.simmap function in phytools (v0.7-70, R package) (35) was used to perform stochastic source mapping based on ARD model and the tip states on the tree, with 10000 generations of MCMC sampling every 100 generations.

### pPCP1 and *pla* analysis

Samtools depth command was used to count the depth of whole pPCP1 plasmid and *pla*-gene for each sample from bam files. The packages of ggplot2 and ggsignif were used to obtain the boxplots and group-wise comparisons (Wilcoxon-test) of the ratio between the depth of *pla* and that of whole pPCP1 plasmid among three subclades of the Second Pandemic. The coverage plots of pPCP1 in CHE1 (Fig. 3) was visualized using Integrative Genomics Viewer(IGV, v2.8.11) (36).

## Data availability

Publicly available genomes are listed in Dataset S1 and Dataset S2.

## Authors’ contributions

B.B. conceived the work; N.C.S. established the author team; B.B., Y.C. and N.C.S. designed research; Y.W. analyzed the data, designed and generated the phylogeny; B.B. wrote the paper with contributions from Y.W., R.Y., Y.C. and N.C.S.; R.Y. and N.C.S. supervised the work.

## Acknowledgement

Katharine Rose Dean is thanked for constructive comments on the manuscript, including improving the language. Two anonymous reviewers on an earlier version of this paper are greatly thanked for comments and suggestions which helped us to improve the paper.

---

**Fig. S1.**
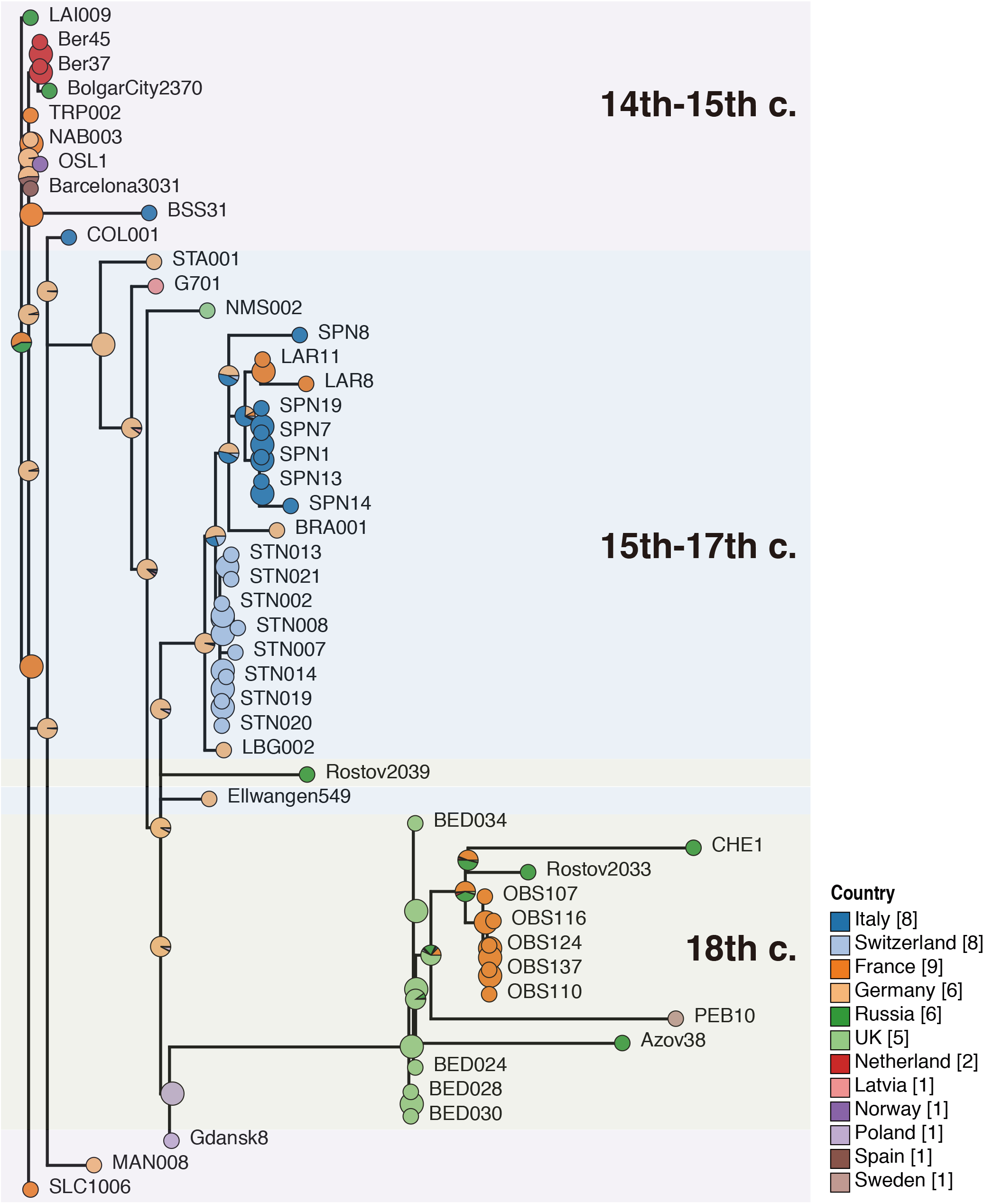
Accessed transmission among different regions. The phylogenetic tree
was obtained using Bayesian MCMC method implemented by phytools package
for R. The pie chart at the internal node of the MCMC tree indicated the ancestor
source probability of its descendent clades. Except for the Gdansk8 node, all
other internal nodes of the tree were sourced from Western Europe, including the
node of 14th-15th c., which appears to have originated in France or Germany with
a high posterior probability. See the text for an explanation.

